# Development of RT-RPA-based point-of-care tests for epidemic arthritogenic alphaviruses

**DOI:** 10.1101/2024.05.14.594209

**Authors:** Sainetra Sridhar, Prince Baffour Tonto, Lily Lumkong, Eduardo Martins Netto, Carlos Brites, Wei-Kung Wang, Bobby Brooke Herrera

## Abstract

Chikungunya (CHIKV), o’nyong-nyong (ONNV), and Mayaro (MAYV) viruses are transmitted by mosquitoes and known to cause a debilitating arthritogenic syndrome. These alphaviruses have emerged and re-emerged, leading to outbreaks in tropical and subtropical regions of Asia, South America, and Africa. Despite their prevalence, there persists a critical gap in the availability of sensitive and virus-specific point-of-care (POC) diagnostics. Traditional immunoglobulin-based tests such as enzyme-linked immunosorbent assay (ELISAs) often yield cross-reactive results due to the close genetic relationship between these viruses. Molecular diagnostics such as quantitative polymerase chain reaction (qPCR) offer high sensitivity but are limited by the need for specialized laboratory equipment. Recombinase polymerase amplification (RPA), an isothermal amplification method, is a promising alternative to qPCR, providing rapid results with minimal equipment requirements. Here, we report the development and validation of three virus-specific RPA-based POC tests for CHIKV, ONNV, and MAYV. These tests demonstrated both speed and sensitivity, capable of detecting 10 viral copies within 20 minutes of amplification, without exhibiting cross-reactivity. Furthermore, we evaluated the clinical potential of these tests using serum and tissue samples from CHIKV, ONNV, and MAYV-infected mice, as well as CHIKV-infected human patients. We demonstrate that the RPA amplicons derived from the patient samples can be sequenced, enabling cost-effective molecular epidemiological studies. Our findings highlight the significance of these rapid and specific POC diagnostics in improving the early detection and management of these arboviral infections.

## Introduction

Chikungunya (CHIKV), o’nyong-nyong (ONNV) and Mayaro (MAYV) viruses are three closely related mosquito-borne viruses that belong to genus *Alphavirus* in the family *Togaviridae*^1^. Their genomes consist of approximately 12 kilobases of positive-sense RNA encased within a nucleocapsid, encoding the four nonstructural proteins (NSP1-4) and five structural proteins including the capsid, three envelope proteins (E1-3), and 6K, across two open reading frames^2^. While evidence of alphavirus-induced human diseases dates back centuries, CHIKV, ONNV, and MAYV were formally isolated in the 1950s: CHIKV in Tanzania (1952), ONNV in Uganda (1959), and MAYV in Trinidad (1954) ^3–5^.

CHIKV is primarily transmitted by *Aedes* mosquitoes^1,3^. Following its isolation in 1952 CHIKV caused several intermittent outbreaks from the 1960s-1980s in the Sub-Saharan Africa, Southeast and South Asia, until its reemergence in 2004, in Kenya^6–9^. It subsequently spread to several nearby East African countries, islands in the Indian ocean, and countries in South/Southeast Asia, resulting in millions of cases^6–9^. Then, in 2011, CHIKV reached the western hemisphere and rapidly diffused through the Caribbean and finally reached South America in 2014, infecting hundreds of thousands of people along the way ^6–8^. Since then, there have been periodic CHIKV outbreaks in all of these regions, as well as some autochthonous cases in parts of Europe, Middle East, and North America^3,6,7^. Overall, according to the European Center for Disease Prevention and Control, CHIKV lays claim to over 100 countries, and in 2024 alone, as of March 31, 160,000 new cases and 50 deaths have been reported^10^.

ONNV, closely related to CHIKV, is limited in its endemicity within Africa, although, reports of the virus reaching countries outside of Africa via international travelers has been previously reported^5,11,12^. Transmitted by *Anopheles* mosquitoes, ONNV has been implicated in two major outbreaks^1^. The first outbreak, spanning 1959-1962, involved countries in East and West Africa, infecting over 2 million people^5,13^. The second outbreak between 1996-1997, emerged in Uganda and then spread to Kenya and Tanzania, with infection rates ranging from 45-68%^1,5^. Since then, isolated cases and smaller outbreaks have been reported across East and West Africa, as well as in travelers returning from those regions^12,13^.

Like ONNV, MAYV, is also geographically limited in its spread, causing sporadic outbreaks in Central and South America, since it was first isolated in 1954 in Trinidad^14,15^. Outside of these, wide ranging seroprevalence (21% to 72%) has been reported all across the region, and sporadic cases occur both within the region and abroad, in Europe and the USA through travelers^14,16^. Although primarily transmitted by *Hemagogous* mosquitoes, certain strains of MAYV have the ability to adapt to replicate in and be transmitted by different, more anthropophilic mosquito vectors^14,17,18^. This feature is a significant public health concern as it has enabled the broader dissemination of related viruses like CHIKV, suggesting a potential global spread of MAYV^14,17,18^.

Infection with these viruses typically results in a febrile illness characterized by fever, mild to severe arthralgia, occasionally accompanied by edema, and/or a rash, but can also lead to a debilitating, long-term arthritogenic syndrome^3^. In the case of CHIKV, approximately 40% of infections progress to chronic arthritis, persisting for up to 3 months. Long-term complications have not been reported for ONNV, whereas, rare chronic arthritis and encephalitic disease has been reported for MAYV^3,19^. Although seldom fatal, the illnesses induced by these viruses substantially impact quality of life. Given the millions of reported cases worldwide, CHIKV, ONNV, and MAYV pose a significant global health risk^1,20^.

The prevalence of these viruses is anticipated to escalate due to the influences of climate change, urbanization, and international travel^20^, necessitating robust surveillance systems to prevent major outbreaks through effective, point-of-care (POC) diagnostic tools. Diagnosis of CHIKV, ONNV, and MAYV infections primarily relies on laboratory testing due to clinical presentations of these viruses overlapping with that of other arboviral diseases like dengue and Zika fever ^1,3,20^. Additionally, the co-circulation of CHIKV and ONNV in Africa, and CHIKV and MAYV in South America, further complicates accurate diagnosis^1,3,20^. Current diagnostic tests are predominantly either serological or molecular techniques, based on the detection of patient antiviral immunoglobulins (IgM and IgG) or viral RNA, respectively^1^.

Commercially available serological tests, such as enzyme-linked immunosorbent assays (ELISAs) detect anti-viral IgM and IgG^1^. IgM antibodies can usually be detected in patient serum or plasma within 6 days after infection and persist for 3-4 months after infection^1^. IgG antibodies are produced subsequently to IgM antibodies and can be detected decades after the initial infection^1,21,22^. ELISAs, however, are reported to have low sensitivity and lack specificity since antibodies tend to cross-react between the three viruses due to their genetic similarity^1^. Plaque reduction neutralization tests (PRNTs) are typically considered gold standard assays to distinguish viral infections based on neutralizing antibodies, but these tests often require enhanced biosafety laboratories and expert handling^23^. Moreover, there is evidence suggesting considerable cross-reactivity between CHIKV and ONNV in PRNTs as well^1,24–27^.

Molecular diagnostics, on the other hand, are more sensitive and specific. Of these, reverse transcription (RT)-PCRs or qRT-PCRs are the most popular^1,20^. There are several commercial and/or in-house RT-PCR tests for CHIKV, ONNV, and MAYV^1,20,28,29^. These assays are reported to reliably detect viruses particularly during the acute phase of the infection (the first 3-4 days), given the high viremia (10^4^-10^8^ RNA copies/mL in the blood) during that time^1^. However, these assays fall short of being ideal POC diagnostics, especially in more resource-limited settings, due to their reliance on expensive and high precision equipment such as thermal cyclers.

Therefore, rapid, inexpensive, sensitive, and specific POC diagnostics are still an unmet need with these three viruses. One possible solution is using isothermal amplification methods to detect viral RNA. These techniques have emerged as faster and less expensive alternatives to PCR, particularly because they do not require precise thermal cycling, and have shorter reaction times^30^. Of these, recombinase polymerase amplification (RPA) is of interest due to its non-complicated assay design, shorter reaction running times, and its potential to be multiplexed^30^. RPA was first described in 2006 and it works by using a single pair of single-stranded DNA primers, recombinase proteins (T4 Uvs X and UvsY), single-strand binding proteins (Gp32), and a strand displacing polymerase (B*. subtilis* polymerase large fragment), to isothermally amplify a double-stranded DNA template in the presence of ATP and magnesium acetate^31^. Reactions are typically run at 37°C-42°C for 5-20 minutes^30^. It can be used in the detection of RNA with the addition of a reverse transcription step, either as a separate step or in a one-step reaction^30,32–34^. Furthermore, the assay can be configured to comply with several detection modalities, including agarose gel electrophoresis, colorimetric readouts, and rapid test/lateral flow assay (LFA) strips^30–35^.

Currently, RPA tests for the detection of these alpha viruses are limited^20^, with the only example being a CHIKV RPA test designed by Patel et al., which was sensitive but cross-reactive with ONNV^36^. In this study, we report the development of three virus-specific and sensitive RPA-based diagnostic tests for CHIKV, ONNV, and MAYV. We initially performed multiple sequence alignments (MSAs) to identify genomic targets for primer design, followed by an extensive primer screen, and then finally re-configuring the successful primer pairs to make the assay compatible with LFA-based detection. The assays target E2 for CHIKV, and NSP2 for both MAYV and ONNV and all three tests can detect down to 10 viral copies on LFA strips. The RPA-based rapid tests were further evaluated on serum and tissue samples from clinically relevant mouse infection models for each virus. Lastly, we applied the CHIKV diagnostic on serum samples from a cohort of patients to demonstrate its clinical potential. Moreover, we were able sequence the RPA amplicons from the patient samples and were able to use the sequences to extrapolate phylogenetic relationships of the isolates. Importantly, we show that, in addition to being sensitive and specific POC diagnostic tools, RPA reactions can also be sequenced, thus enabling more inexpensive molecular epidemiological efforts to better characterize these viruses and monitor outbreaks.

## Results

In order to obtain primer pairs and probes capable of detecting RNA from all strains of a given virus, we wanted to target them to the most conserved regions of that virus’ genome. To identify such targets for primer design, we used BLAST to construct MSAs of a set of complete genomes for each virus. We then analyzed the consensus sequence generated from the MSA to identify the most conserved genomic regions, focusing our analysis on the 4 NSPs, capsid, E1 and E2. The two remaining structural proteins—6K, E3—were overlooked due to their smaller size and therefore less area within which to design a set of primers. The top 4 most conserved genomic regions identified for each virus were as follows: NSP2, NSP4, E2 and E1 for CHIKV; NSP1, NSP2, NSP4, E2 for ONNV; NSP1, NSP2, NSP4 and E1 for MAYV (Fig. 1A-C). Primers were designed within the identified genes for each virus, with one exception: for ONNV, we opted to design primers within E1 rather than NSP4. This was due to its high degree of homology, with less than 40% sequence divergence within the Semliki Forest Serocomplex, the antigenic complex to which the CHIKV, ONNV, and MAYV belong and also due to the high utility of E1 as a target in existing molecular assays for these viruses^1,37^.

**Figure 1:**
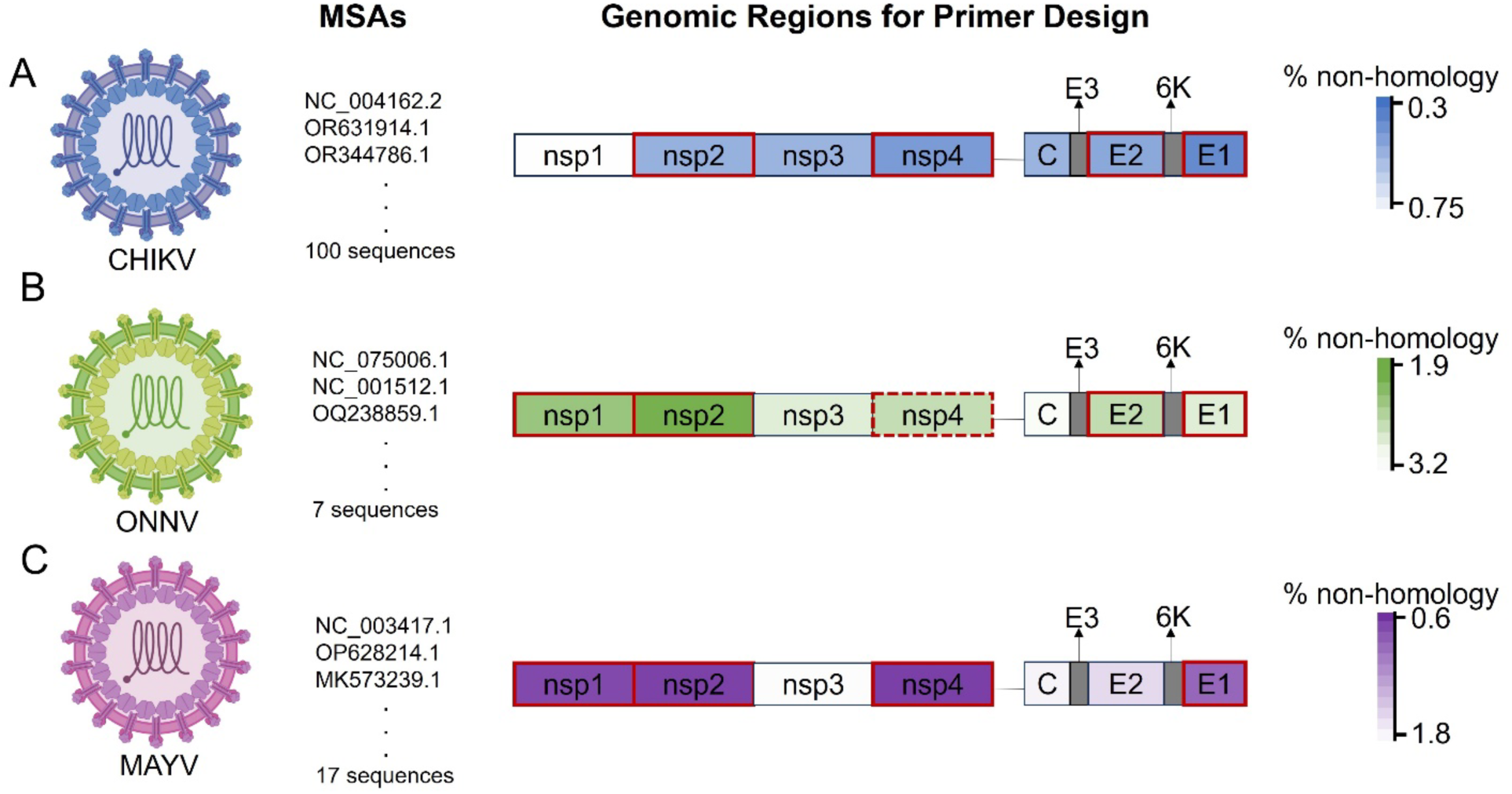
Schematic depicting the regions selected for primer design. Multiple sequence alignments (MSAs) were assembled for (A) CHIKV, (B) ONNV and (C) MAYV using the indicated number of sequences following which the consensus sequence was analyzed focusing on the four NSPs, the Capsid, E2 and E1 to identify the most conserved regions for each virus. The heatmaps show the percent non-homology calculated for each region and the top four most homologous sites (with the darkest shading), highlighted in a red border, were selected for primer design. For ONNV, the analysis identified NSP4, highlighted with a discontinuous border, was identified as the fourth target for primer design, however, we chose to prioritize designing primers in E1 instead.

16 primer pairs were designed for ONNV and 17 for CHIKV and MAYV, spanning the genomic regions identified through the MSAs. It was ensured that the primers followed the requirements for RPA primer design: a length of 30-35 nt, a GC content of 40-60% and absence of palindromes or tandem nucleotide repeats^32,35,38^. Additionally, the designed primers were mapped onto the genomes of the other two viruses, to ensure sufficient mismatches were present to minimize chances of cross-reactivity^38^. To screen the designed primers, viral genomic RNA (gRNA) was first reverse transcribed and then an equivalent of 10^5^ viral copies in complementary DNA (cDNA) was used as input into each RPA reaction. The reactions were run for 20 minutes, at three temperatures— 37°C, 39°C and 41°C—spanning the typical RPA temperature range^38^. They were evaluated by running the products out on an agarose gel (Fig. 2A-C). Each primer pair varied in its ability to amplify cDNA, and this activity sometimes influenced by temperature, for example: at 37°C, CHIKV primers #7, 9, 11 fared better than #3, 6, 10, and #7 fared better at 37°C than at 39°C (Fig. 2A). At the end of the screening, several successful primer-pair/reaction-temperature combinations were identified, and the following combinations were selected: #9 at 37°C was identified for CHIKV, #2 at 41°C for ONNV and #1 at 39°C for MAYV. Their respective targets are within the E2 gene for CHIKV and within NSP2 for both ONNV and MAYV. The analytical sensitivity of these three primer-pairs was evaluated by using a serial dilution of cDNA, ranging from 10^6^ viral copies to 1 viral copy. Also included were reactions using 10^5^ viral copies of the other two viruses to evaluate cross-reactivity of the primers. The limits of detection (LODs) for these three primers, processed via agarose gel electrophoresis, was found to be approximately 10^2^ viral copies for CHIKV and ONNV and between 10^3^-10^2^ viral copies for MAYV, and no cross-reactivity to cDNA from the other two viruses was seen (Fig. 3).

**Figure 2:**
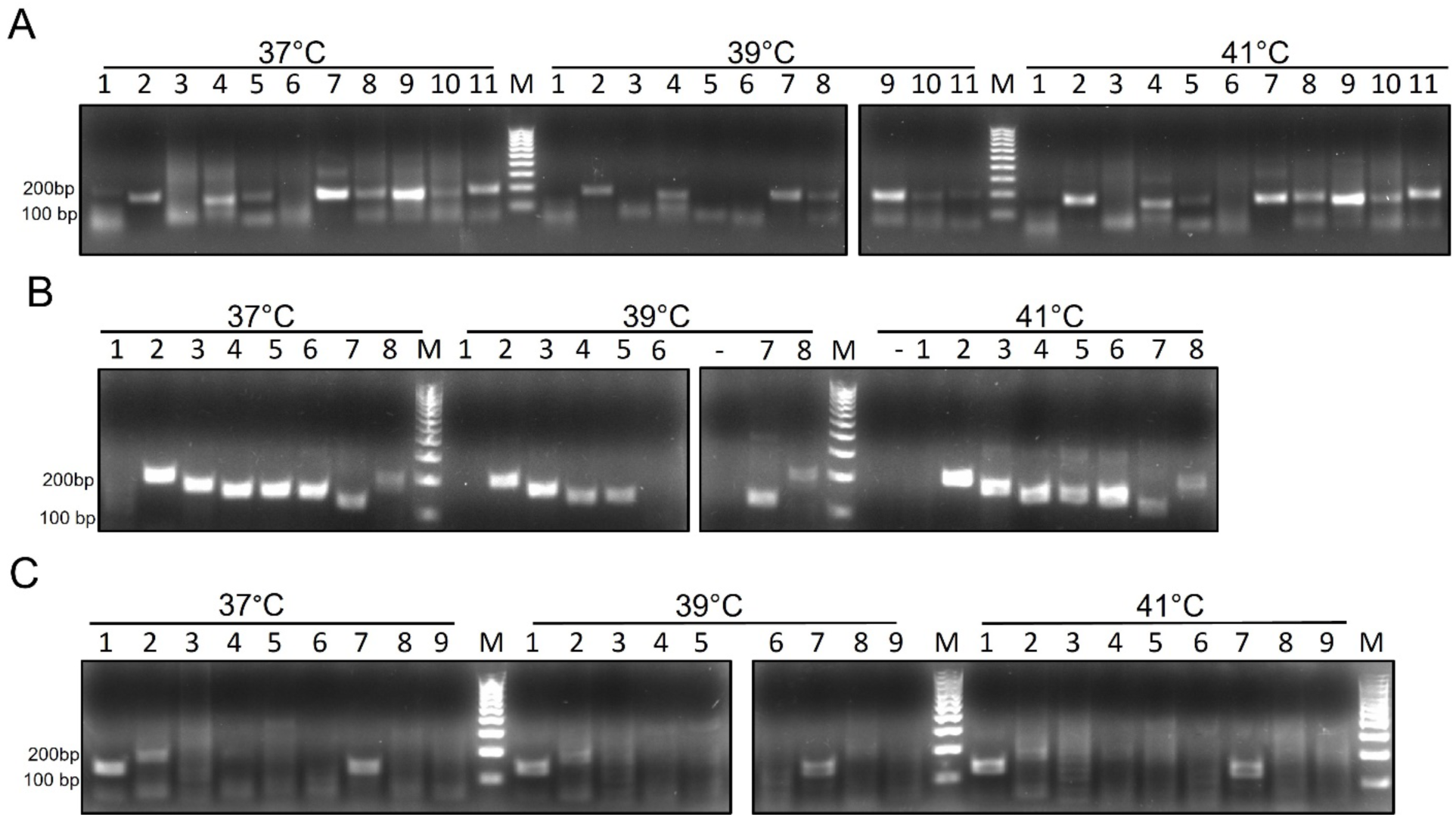
Primer screens conducted by agarose gel electrophoresis of RPA products. All 17 primers for CHIKV, 16 primers for ONNV and 17 primes for MAYV were screened in 20-minute RPA reactions using 10^5^ viral copies with cDNA as input, The reactions were run at three temperatures as indicted. Depicted here is a representative subset of those screened primers, (A) corresponding to CHIKV, (B) to ONNV, and (C) to MAYV.

**Figure 3:**
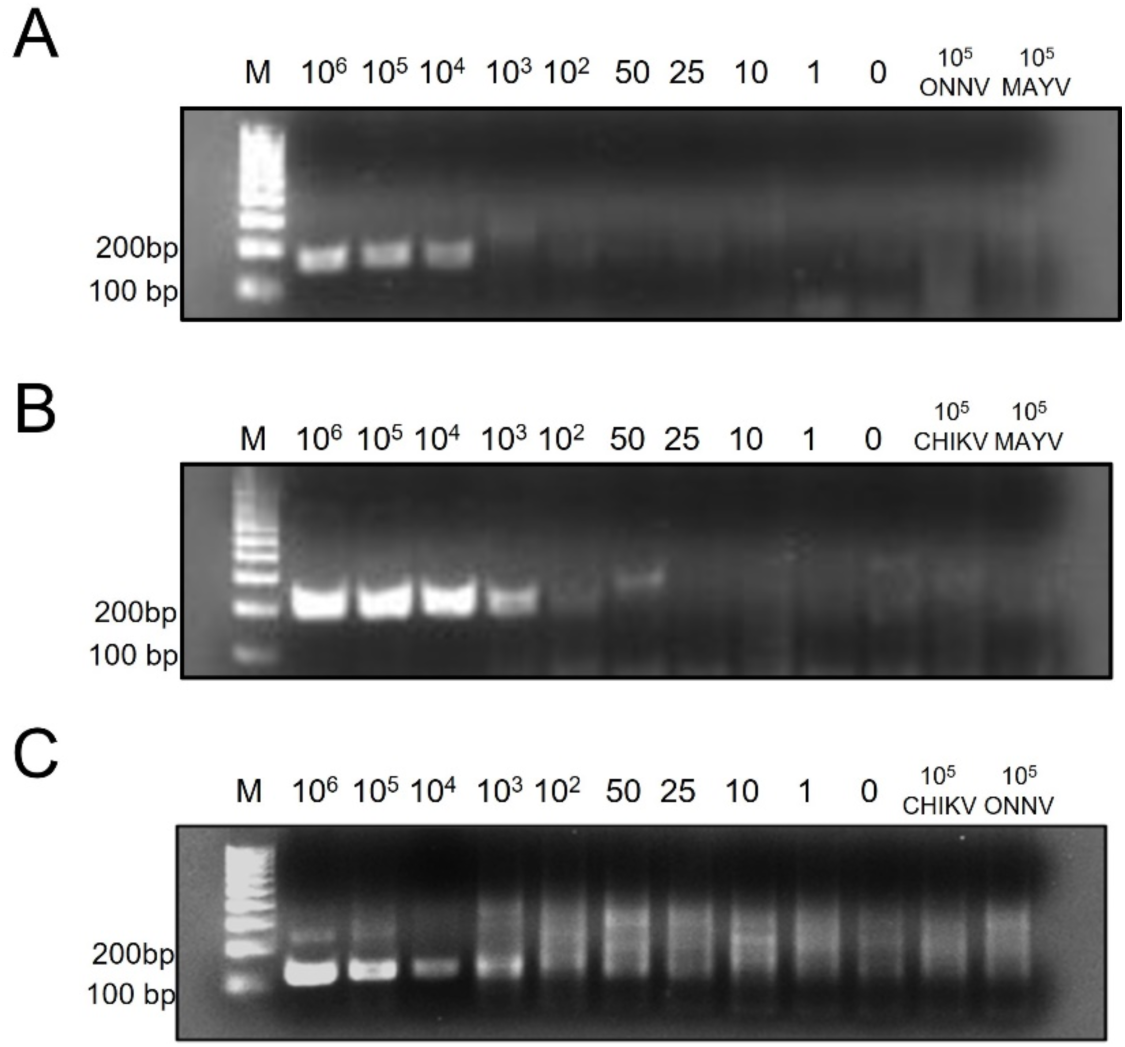
Analytical sensitivities of the designed primers on an agarose gel. Candidate (A) CHIKV, (B) ONNV, and (C) MAYV primer pairs identified in the screen were evaluated on a serial dilution of cDNA inputs ranging from 10^6^ viral copes to 1 viral copy. Cross reactivity of these primers was also tested using 10^5^ copies of cDNA from the other two viruses. The reaction inputs for each well are denoted in the image. The sensitivity was determined to be 10^2^ for CHIKV and ONNV, and between 10^3^ and 10^2^ for MAYV, with not cross-reactivity.

Since we sought to design rapid tests for these three viruses, our next step was to re-configure our assays to enable the detection of the reaction products on LFA strips. For this, two approaches were used: hybridization and nfo-probes. At the outset, both approaches involve replacing one of the reaction primers with a 5’labelled version to generate a singly labelled amplification product. The first method involves a separate hybridization step, where the singly labelled amplification product, is denatured and then hybridized with a labelled probe^34,39^. The second approach involves introducing an nfo-probe, which are labelled probes with an internal abasic site, into the reaction, along with an endonuclease (nfo or Endo IV) that cleaves at that abasic site upon probe hybridization^31^. Ultimately, both approaches result in an amplification product that is doubly labelled—one label with affinity for test line on the strip, and the other, with affinity for the antibody conjugated to the gold nanoparticles (Fig. 4A). The LFA strips we used had two test-bands, one coated with streptavidin and the other with an anti-digitonin antibody. The gold nanoparticles in the strip are conjugated to an anti-FITC antibody. We therefore opted to label the reverse primers using either 5’biotin tag (for CHIKV) or 5’Digitonin tag (for ONNV and MAYV) and labelled all the probes with a 5’FAM tag.

**Figure 4:**
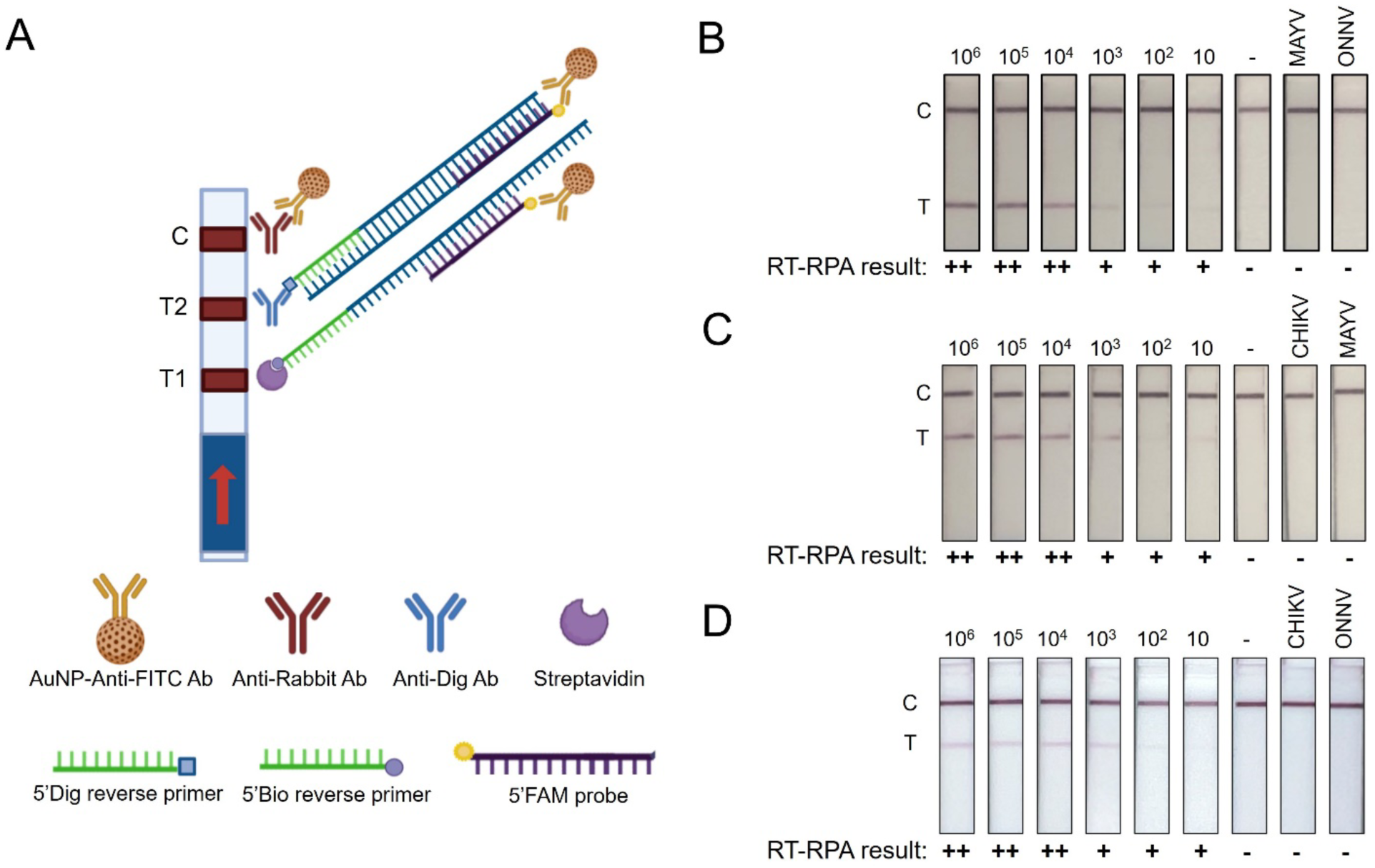
Detecting RPA reactions on LFA strips. A) Schematic representation of the Millenia Biosciences 2T test strips utilized in this study. The strips have two test bands “T1” and “T2,” one coated with streptavidin and the other with an anti-Digitonin antibody. The gold nanoparticles (AuNPs) are conjugated to an anti-FITC antibody. Accordingly, all our reverse primers were either tagged with a 5’Biotin tag (CHIKV) or a 5’ Digitonin tag (ONNV and MAYV), and their probes-hybridization probe for CHIKV and MAYV or nfo-probe for ONNV-were all 5’FAM tagged. Shown in the figure are depictions of the reaction product from the CHIKV assay, that uses a hybridization probe, binding at T1 and the ONNV assay, that uses a nfo-probe, binding at T2. The MAYV RPA product would appear like the CHIKV product since it also uses hybridization probes, however, it would bind at T2, since its reverse primer has a 5’digitonin tag. (B-D) show the analytical sensitivity of the rapid-test versions of the CHIKV (B), ONNV, (C) and MAYV (D) assays. The reactions used a serial dilution of cDNA as their input. Also included are reactions with 10^5^ copies of cDNA from the other two viruses to assess cross-reactivity. RPA reaction products were diluted and applied to the strips and then immersed in the manufacturer-provided assay buffer for up to 30 minutes. The strips were photographed using a smartphone camera. All three assays were able to detect down to 10 copies of viral cDNA and were not cross-reactive.

We chose the hybridization option for the CHIKV and MAYV tests, and the nfo-probe option for ONNV. The reaction temperatures were maintained at the same values that were identified in the agarose gel-based primer screen for the ONNV and MAYV tests. However, for CHIKV, running the reaction at 39°C instead of 37°C resulted in darker coloration of the test-band on the LFA strip, prompting us to modify the running conditions accordingly. Sensitivity of the rapid test was determined in the same manner as described for Figure 3, using a serial dilution of viral cDNA. All three tests were highly sensitive, with the ability to detect down to 10 viral copies, and similarly to their un-labeled counterparts (Fig. 3), they did not cross-react with 10^5^ cDNA copies from the other two viruses (Fig. 4B-D). Furthermore, we conducted a broader cross-reactivity screen for these three rapid tests using cDNA from other arboviruses and saw no cross reactivity, enabling us to confirm that the designed assays were highly specific to their intended targets (Supplementary Table 3).

Having designed sensitive and specific RT-RPA-based rapid tests for CHIKV, ONNV, and MAYV, we then sought to evaluate their performance on clinically relevant samples. For this, adult B6 mice homozygous for interferon alpha/beta/gamma receptor (IFNRα/β/γ) knockouts, were inoculated with either CHIKV, ONNV, or MAYV at a dose of 1000 PFU via subcutaneous footpad injection. RNA was then extracted from serum collected 1- and 3-days post-infection (dpi) and from the spleens, livers, and kidneys. The extracted RNA was subjected to the designed RT-RPA tests and also to RT-PCR for comparison (Fig. 5A-C). The CHIKV test was able to detect virus in 2 out of 3 of the 1-dpi serum samples, 3-dpi serum samples, and livers. But all three kidney and spleen samples produced a positive test. The CHIKV RT-PCR test in comparison only detected viral RNA in two samples: one of the 3-dpi serum samples and in one of the livers (Fig. 5A). The ONNV rapid test detected viral RNA in 1 out of 2 of the 3-dpi serum samples, spleens, and kidneys and both liver samples (Fig. 5B). Several of the RT-RPA-negative ONNV samples did however test positive with RT-PCR. All but one of the MAYV samples tested positive on both RT-RPA and RT-PCR, with the only negative result coming from one of the 1-dpi serum samples (Fig. 5C).

**Figure 5:**
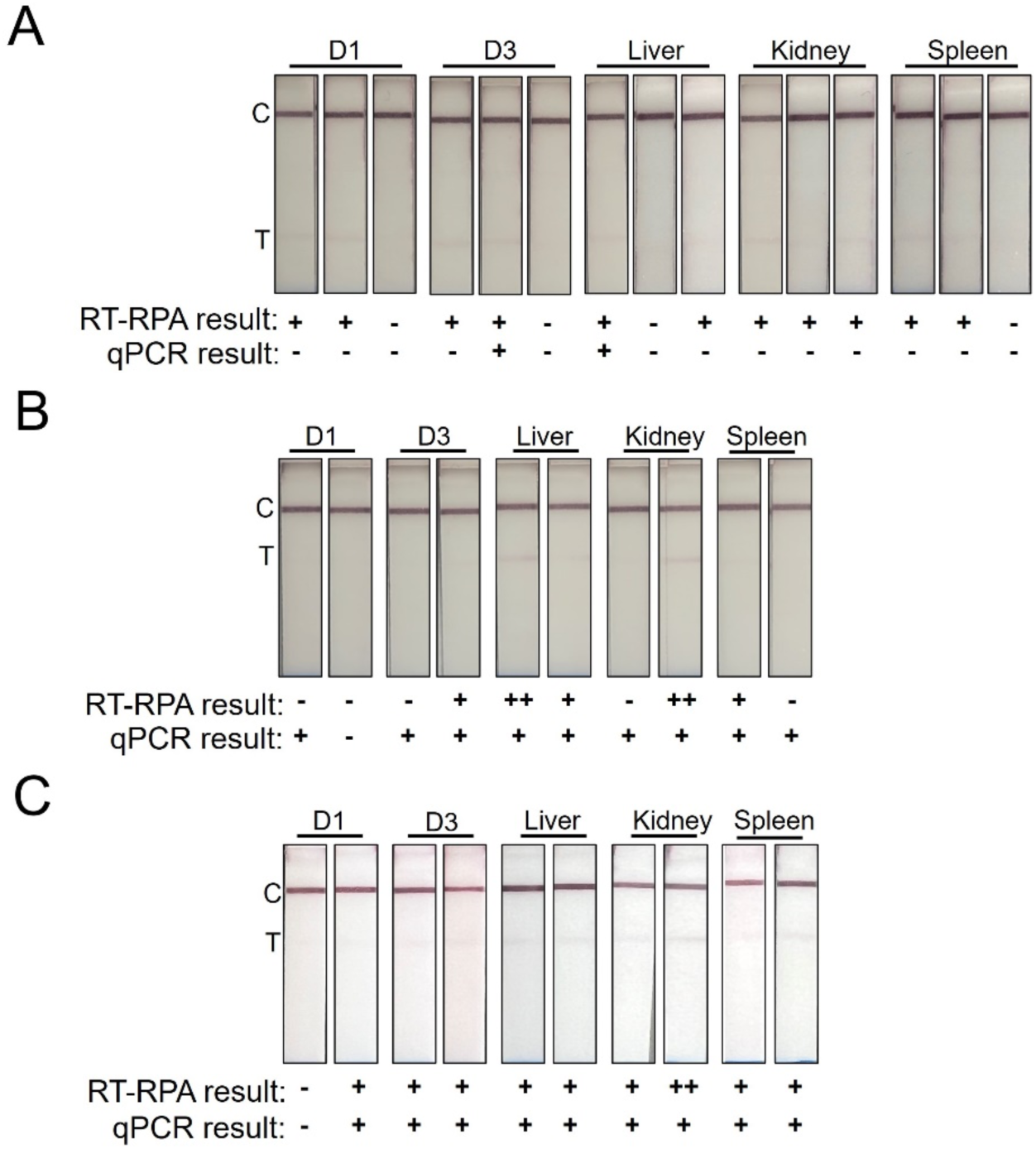
Detecting alphaviruses in serum and tissues from mouse infection models using RT-RPA rapid tests. IFNRα/β/γ -/- mice were infected with 1000 PFU of (A) CHIKV (181/25 strain, n=3), (B) ONNV (NR-50081 strain, n=2) and (C) MAYV (NR-51661 strain, n=2) via subcutaneous footpad injection. Serum was collected 1- and 3-days post infection, and organs were harvested at day 3 for CHIKV and ONNV and at day 5 for MAYV, when the mice experienced mortality. RNA was extracted, reverse-transcribed and subjected to RPA using labelled primers. The reaction products were applied to the test strips and were photographed using a smartphone camera. The extracted RNA from each sample was also subjected to RT-PCR for comparison. The results of both assays for each sample are indicated with + (or ++) and –, as observed.

As a final evaluation of the clinical potential of our assays, we applied our CHIKV RT-RPA rapid test to 5 patient serum samples from Brazil that tested positive for CHIKV by RT-PCR. Also included in this analysis were Zika- (ZIKV) and dengue- (DENV) RT-PCR positive samples, as negative controls for the CHIKV assay. All CHIKV RT-PCR positive patient samples were also positive on our rapid test, whereas the ZIKV and DENV RT-PCR positive samples tested negative on our rapid test (Fig. 6A). We also were able to subject the RPA amplicons from the 5 CHIKV positive samples to sanger sequencing and the resulting sequences spanned about 65% of the target amplicon in the CHIKV E2. We generated a phylogenetic tree using aligned sequences of those derived from 5 patients, the product from an RPA reaction using the same cDNA as the tests/screens in Figures 2-4 (from the 181/25 strain), and the E2 sequences of various isolates and strains (Fig. 6B). We found that the five patient sequences clustered close together and two of them clustered close to another known CHIKV Brazilian isolate with an East-Central-South-African (ECSA) genotype. The patient sequences, the Brazilian-ECSA isolate, and another African strain all clustered separately from a CHIKV Brazilian isolate with an Asian genotype, and the CHIKV 181/25 strain (including the 181/25 sequence obtained from our RPA reaction) which is a known live-attenuated derivative of a Southeast Asian CHIKV strain^40^.

**Figure 6:**
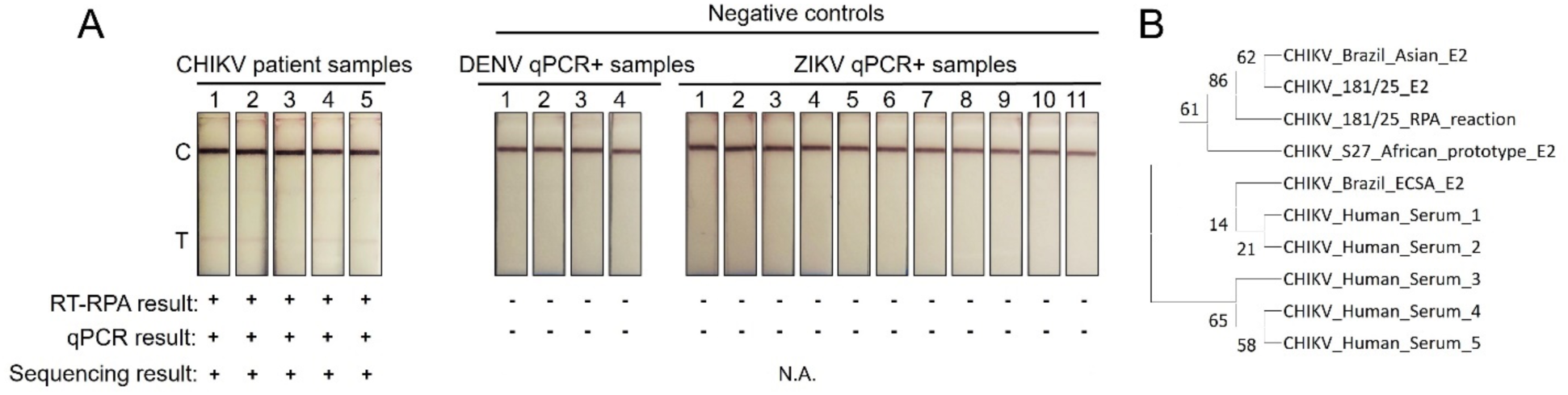
Detecting CHIKV from patient samples using the CHIKV RT-RPA test. A) 5 CHIKV qPCR-positive patient serum samples from Brazil were evaluated using our designed RT-RPA test, with Zika and dengue qPCR-positive patient sera as negative controls. RNA was extracted and reverse transcribed, and the resulting cDNA was subjected to RPA. The results are as indicated in the figure with either + for a positive result or - for a negative result. Additionally, the RT-RPA amplicons from the 5 patient samples were subjected to sanger sequencing and the resulting sequence corresponded with the intended target region in E2. B) Phylogenetic tree constructed from aligning the sequences obtained from sanger sequencing of the 5 patient RPA amplicons (CHIKV_Human_Serum1-5) and RPA product from the same gRNA used in the screens and initial tests (181/25_RPA reaction) with the E2 sequences of various CHIKV strains and isolates. The accession numbers of the strains included in this analysis are as follows: OQ148623.1 (CHIKV_Brazil_Asian, a Brazilian CHIKV isolate with an Asian genotype), L3766.1 (CHIKV_181/25, a live attenuated derivative of an Asian strain of CHIKV, and the source of the gRNA used in the screens and initial tests of our RT-RPA assay), NC_004162.2 (CHIKV_S27_African_prototype, an African strain of CHIKV, and the genome that was used as the basis for primer design in this study), KU940225.1 (CHIKV_Brazil_ECSA, a Brazilian CHIKV isolate with an East-Central-Southern-African genotype). A ClustalW alignment of the sequences was used to construct a maximum-likelihood tree based on the Tamura-Nei model and 500 bootstrapping replications using Mega11.

## Discussion

In this study, we describe the development of three RT-RPA based rapid tests for the detection of CHIKV, ONNV and MAYV. The three viruses are closely related alphaviruses, belonging to the same serocomplex, with ONNV and CHIKV being the closest related viruses in that group^37^. Currently, there is a lack of sensitive and specific POC diagnostic tools for these three viruses, given the cross reactivity of common serological tests, and the expensive cost of running common molecular diagnostic tests like RT-PCRs^1,20^. We sought to bridge that gap by designing RT-RPA based rapid tests, that utilize the isothermal amplification technique, RPA, and are detected on rapid test strips, making them ideal for use at the POC. We show that our three tests were sensitive, with the ability to detect down to 10 viral copies, and specific, with no cross reactivity to related alphaviruses or other arboviruses.

We further demonstrated that our tests are capable of detecting viral RNA from more clinically relevant samples, starting with those obtained from mouse infection models. Mice deficient in interferon signaling are commonly used to model CHIKV, ONNV, and MAYV infections because wildtype mice are not as susceptible and are able to rapidly clear infections from these viruses^41,42^. In our mouse experiments, we were able to detect viral RNA in the day 3 sera more reliably than in the day 1 sera, consistent with the trend of viremia peaking around day 2-3 for arboviral infections in this mouse model^42^. We were also able to detect viral RNA in the spleens and livers, which, along with joints and muscles, are considered common replication sites for alphaviruses in their hosts, allowing for their efficient dissemination thereafter^41,43–45^. We were also able to detect viral RNA in the kidneys of infected mice, and while they not a common target or replication site for alphavirus infections, viral loads have been detected in the kidneys of interferon signaling-deficient mice infected with100 PFU of CHIKV^46^. Additionally, we were only able to produce faint positive test bands on our rapid-test strips for most of these samples. One possible explanation for this could be that these mice succumbed to the infections quite rapidly, and possibly before the viruses had a chance to replicate to produce robust viral loads. It is worth noting that the 3 CHIKV infected mice and the 2 ONNV infected mice experienced mortality at day 3, and the 2 MAYV infected mice succumbed to the infection at day 5. This is consistent with what with what is known about the behavior of CHIKV, ONNV, and MAYV infections in similar mouse models^41,42,45,47–52^.

Lastly, applying the CHIKV test to patient samples enabled us to further establish the clinical potential of our RT-RPA tests. The test reliably produced a positive result for all of the known CHIKV positive samples tested, and we were able to also confirm this result by subjecting the RPA product to sanger sequencing. The ability to sequence RPA reaction products has been previously demonstrated, using a range of sequencing platforms from Illumina to MinION Nanopore^53–55^. We subjected our RPA products to same-day sanger sequencing and were subsequently able to take it a step further and conduct a phylogenetic analysis of the obtained sequences with success. Short amplicon sequencing from PCR products to characterize the phylogenetics of viral isolates has been done before^54^, and while it is important to acknowledge that short amplicon sequencing is limited in its ability to fully characterize viral evolutionary dynamics, the ability of RPA products to be analyzed in a similar manner opens up the possibility of conducting molecular epidemiological studies in a more rapid and inexpensive manner, which would supplement diagnostic efforts and help better monitor outbreaks.

One improvement that would further bolster these assays would be optimizing them to function as one-step RT-RPA reactions. Secondly, we had designed the MAYV and ONNV assays to produce products that have a different tag than the CHIKV assay products (i.e., digitonin rather than biotin) because we had initially aimed at multiplexing their detection on our double test-band LFA strips. Further optimization to enable this would allow for the creation of two alphavirus tests, one for the South American viruses-CHIKV and MAYV, and the other for the African viruses-CHIKV and ONNV. Lastly, being able to detect RNA directly from crude biological samples such as plasma/serum, urine, or saliva, without an RNA extraction step, would truly enable field deployment. This is possible and has been attempted before, because compared to PCR, RPA is less susceptible to inhibitors present in crude samples^35,56^.

Despite their spread and disease severity, approved vaccines and antiviral therapies for these three alphaviruses are limited, with the first approved CHIKV vaccine being as recent as of November 2023^57^. As these efforts ramp up with the increasing recognition of the threats posed by these viruses, effective POC diagnostic tools will be critical for the efficient monitoring of outbreaks, and for the dissemination of prophylactics and therapeutics. Given their potential as sensitive, specific, POC diagnostics, we think that the RT-RPA assays described here are a promising step forward towards that goal.

## Methods

### Selecting regions for primer/probe design

MSAs were constructed using NCBI BLAST by first querying a complete genome for each virus (NC_004162.2 for CHIKV, NC_075006.1 for ONNV, NC_003417.1 for MAYV). The search results were filtered down to obtain a set of complete genomes with over 90% similarity to the query sequence. Those sequences were aligned using BLAST’s MSA tool and viewed using the NCBI MSA viewer 1.25. The MSA viewer was used to generate a consensus sequence with a 70% threshold, i.e., if >=70% of the aligned sequences have the same nucleotide at the same position in the alignment, the consensus sequence will reflect that, and, if there is less than 70% consensus at a particular position, an ambiguous IUPAC symbol is used instead. The positions with a less than 70% consensus (i.e., with an ambiguous IUPAC symbol in the consensus sequence) were considered “mismatches” while calculating the % non-homology for each gene of interest. Finally, the % mismatch values for the various genes were compared and the top 4 most conserved, i.e., with the lowest % mismatch values were selected for primer design. A list of all the aligned accession numbers is provided in Supplementary Table 1.

### Primer and Probe design

Genomes, NC_004162.2, NC_075006.1 and NC_003417.1 for CHIKV, ONNV and MAYV, were respectively used to design primers and probes, within the genomic regions of interest identified through the MSAs, for each virus. Primers were designed manually, adhering to the following rules: a length of 30-35 nt, a GC content of 40-60% and absence of palindromes or tandem nucleotide repeats^32,35,38^. The target amplicons were between 120-200 bp in size and had a GC content of 40%-60%. Finally, the designed primers were mapped onto another MSA of the complete genomes of the three viruses to ensure that sufficient mismatch was present to prevent cross reactivity^38^. After the primer screen, probes were designed corresponding to the primer pair identified in the screen for each virus. Nfo probe for the ONNV assay was designed as described by Piepenburg et al., ensuring a complete length of about 45 nt, with at least 30 nt preceding the abasic site, and 15 nt following it^31^. The hybridization probes for the CHIKV and MAYV assays were designed as described by Qian et al., to be around 15-25 nt in size, and binding to a region within the amplicon, avoiding overlap with the primer binding sites^33^. The designed primers were synthesized by Integrated DNA Technologies. A list of the RPA primers and probes designed and used in this study are provided in supplementary Table 2.

### RNA extraction

The QiaAmp viral Mini kit (Qiagen) or the PureLink Viral RNA/DNA Mini kit (ThermoFisher) were used for extraction of RNA from serum or culture supernatant for ONNV (NR-51661, BEI Resources Repository), following the manufacturers’ instructions. RNA from mouse tissues was extracted using the RNeasy Lipid tissue Mini Kit (Qiagen) following manufacturer instructions. The concentration of the RNA extracts was determined using a NanoDrop One device (ThermoFisher).

### cDNA synthesis

Viral gRNA for CHIKV (181/25, BEI Resources Repository) and for MAYV (NR-50081, BEI Resources Repository) and extracted RNA from ONNV (NR-51661, BEI Resources Repository) propagated on Vero cells (CCL-81, ATCC) was used. Viral RNA was converted to cDNA using the SuperScript IV Vilo master mix (ThermoFisher), following the manufacturer’s instructions. RNA input of 2x10^7^ viral copies was used in a 20 µL cDNA synthesis reaction to obtain cDNA with a concentration equivalent to 10^6^ copies/µL. For the primer screen, the cDNA stock was diluted to 10^5^ copies/µL and 1 µL was used for the RPA reactions. For the sensitivity assays, the cDNA stock was serially diluted to obtain a range of concentrations, to enable inputs ranging from 10^6^ copies to 1. For the serum and tissue samples, a fixed volume of RNA (8 µL and 2 µL, respectively) was used as input into cDNA synthesis, and either 1µL or 2µL (only for the patient samples) of the cDNA was used as input into RPA.

### RT-RPA reactions using unlabeled primers

RPA reactions were performed using the TwistAmp Basic Lyophilized kit (TwistDx), following the manufacturer’s instructions, but scaling measurements down to 10 µL reactions. Briefly, lyophilized mix was rehydrated using 29.5 µL of the provided rehydration buffer and 10 µL of nuclease free water, and 7.9 µL of the rehydrated mix was used per 10µL reaction, along with 0.3 µM of each primer, 280 mM magnesium acetate, and the requisite amount of cDNA. The reactions were incubated for 20 minutes at either 37°C, 39°C or 41°C, as indicated in the text. Diluted reaction products (1:5) were run on a 2.5% agarose (BioRad) gel in 1x TAE buffer (ThermoScientific) and stained with Gelgreen (Biotium). For the sensitivity assay gels, RPA products were purified using the PureLink PCR purification kit (ThermoFisher), following the manufacturer’s instructions.

### RT-RPA reactions using labelled primers

For the hybridization method, reactions were set up as described above with one modification—the replacement of the reverse primer with a 5’biotin- or 5’digitonin-labelled version, keeping the concentration the same. The hybridization step was performed as described by Qian et al. ^33^. Briefly, after the RPA reaction run, 1 µL of 5 µM hybridization probe (final concentration of 0.167 µM) and 19 µL of hybridization buffer (10 mM Tris HCl at pH 8.5) was added to each reaction. This was heated to 95°C for 3 minutes and then allowed to gradually cool to room temperature to enable hybridization. The nfo-probe reactions were performed as described by Piepenburg et al^31^. The basic master mix was modified, replacing a portion of the water with nfo or Endonuclease IV (New England Biosciences) at a final concentration of 0.2 U/µL and nfo-probe to a final conc of 0.1 µM. The reverse primer was also replaced with a 5’digitonin-labelled version, keeping the concentration the same.

### Detection of RPA products on LFA strips

The products from the hybridization and nfo-probe reactions were processed using the HybriDetect 2T LFA strips (Milenia Biosciences), following the manufacturer’s instructions. Briefly, unpurified reaction products were directly diluted in the provided assay buffer (1:20 for the nfo-probe reactions and 1:15 for hybridization reactions) and applied to the strip. The strip was then immersed in 70 µL of the provided assay buffer and was allowed to react for 5-30 minutes. The strips were imaged using a smartphone camera.

### qRT-PCR

For the CHIKV, ONNV and MAYV qPCRs conducted for this study, the PrimeTime one-step qRT-PCR kit (Integrated DNA Technologies), along with primers and TaqMan probes were used. CT values were recorded using a QuantStudio3 device (ThermoFisher). The primer and probe sequences and the publications they were derived from are as follows:

1. CHIKV- 5’AAGCTCCGCGTCCTTTACCAAG 3’, 5’CCAAATTGTCCGGGTCCTCCT 3’, 5’/FAM/CCATGTCTTCAGCCTGGACACCTTT/TAMRA/3’^48^.
2. ONNV- 5’CGCAGCTTACGGGTTTCATA3’, 5’GCAACGCCTTCAGAAACGC3’, 5’/FAM/TGCTCTACTCTGCATTGCAAGA/BHQ1/3’^29^.
3. MAYV- 5’AAGCTCTTCCTCTGCATTGC3’, 5’TGCTGGAAACGCTCTCTGTA3’, 5’/ CalFluorOrange560/GCCGAGAGCCCGTTTTTAAAATCAC/ BHQ-1/3’^28^.

### Preparation of Mouse Samples

Adult C57BL/6J-congenic Ifngr1 Ifnar1 double mutant mice (JAX stock #029098) were inoculated with 1000 PFU of either CHIKV (181/25 strain, BEI Resources), ONNV (NR-50081, BEI Resources), or MAYV (NR-51661, BEI Resources) via subcutaneous injection on the ventral side of the left footpads. Submandibular blood was collected was collected at day 1 and/or day 3 post infection, and into heparin coated tubes and centrifuged at 2000 x g for 15 minutes, to obtain serum. The mice succumbed to the infections at day 3 (CHIKV and ONNV) and at day 5 (MAYV) at which point, the liver, spleen, kidneys, and cardiac blood were collected. These samples were subjected to RNA extraction and RT-RPA as described above.

### cDNA from arbovirus infected subjects

This study was approved by the Institutional Review Boards (IRBs) of the University of Hawaii (2019-00910) and Federal University of Bahia, Brazil (Comite de Etica em Pesquisa da Maternidade Climerio de Oliveira/UFBA, Brazil, approval number:

25336819.3.0000.5543/4.691.233, 2019). Blood samples from participants with acute febrile illness suspected of DENV, ZIKV or CHIKV infections were collected between 2019 and 2023 in Feira de Santana and Salvador, Brazil. RNA was extracted from serum samples and subjected to a Tetraplex RT-PCT assay, with in-house designed primers and TaqMan probes for DENV, ZIKV, and CHIKV^58^. Human RNase P was also included in the assay as an internal control^59^. Extracted RNA was separately subjected to reverse transcription and the cDNA derived for the DENV, ZIKV, or CHIKV RT-PCR-positive samples was used for RT-RPA reactions as described above.

### Sanger sequencing and phylogenetic analysis

For the amplicons that were sequenced, the TwistAmp Basic Liquid kit (TwistDx) was used instead of the lyophilized kit. 25 µL of the 2x Reaction Buffer, 5 µL of the 10x Basic E-mix, 2.5 µL of the 20x Core Mix, 1.8 mM dNTPs, 6 µL of water were combined and mixed. 7.9 µL of that master mix per 10 µL reaction was then combined with the unlabeled primers, cDNA, and magnesium acetate and processed as described above. RPA amplicons were confirmed by 2.5% agarose gel electrophoresis and subsequently sent to Eton Biosciences along with an aliquot of the forward primer for their same day sequencing service. The sequences that were returned were aligned using ClustalW and a maximum-likelihood phylogenetic tree was constructed on Mega11, using the Tamura-Nei model and 500 bootstrap replications^60,61^.

## Acknowledgements

We would like to thank Rutgers Global Health Institute, Rutgers Robert Wood Johnson Medical School, and The Child Health Institute of New Jersey for their continued support. This work was supported by grants R01AI149502 (WKW) from the National Institute of Allergy and Infectious Diseases, National Institutes of Health (NIH) and 404193/2019-6 (EMN) from the Brazilian National Council for Scientific and Technological Development (CNPq).

## Author contributions

SS designed probes and primers, planned, and carried out most of the described experiments, analyzed experimental data, and wrote the original draft of manuscript. PBT and LL planned and carried out experiments, particularly the mouse infections and RNA extractions. EMN and CB collected acute serum samples from patients and confirmed CHIKV, DENV or ZIKV by RT-qPCR. WKW extracted RNA from those serum samples and synthesized cDNA. BBH conceived of the original idea, supervised the work, and planned and carried out experiments. All authors contributed to the interpretation of the data and played a role in reviewing and editing the manuscript.

## Data Availability Statement

Data produced as part of the current study are available in the manuscript or upon request from the corresponding author.

