## Supplementary Tables for "Development of RT-RPA-based point-of-care tests for epidemic arthritogenic alphaviruses"

Supplementary Table 1. Accession numbers used for CHIKV, ONNV, and MAYV MSAs

| Accession numbers used for the CHIKV MSA | Accession numbers used for the ONNV MSA | Accession numbers used for the MAYVV MSA |
|---|---|---|
| NC_004162.2 | NC_075006.1 | NC_003417.1 |
| OR631914.1 | NC_001512.1 | OP628214.1 |
| OR344786.1 | OQ238859.1 | MK573239.1 |
| OR283254.1 | ON595759.1 | MK573238.1 |
| OQ750694.1 | MF409176.1 | MH513597.1 |
| OQ750692.1 | M20303.1 | KY985361.1 |
| OQ750691.1 | KX771232.1 | KY618135.1 |
| OQ750690.1 | AF079457.1 | KY618133.1 |
| OQ750689.1 | | KY618129.1 |
| OQ750688.1 | | KY618127.1 |
| ON009843.1 | | KX496990.1 |
| MW581885.1 | | KT818520.1 |
| MW581884.1 | | KT754168.1 |
| MW581883.1 | | KP842820.1 |
| MW581882.1 | | KP842819.1 |
| MW581881.1 | | KP842818.1 |
| MW581880.1 | | AF237947.1 |
| MW581879.1 | | |
| MW581878.1 | | |
| MW581877.1 | | |
| MW581876.1 | | |
| MW581875.1 | | |
| MW581874.1 | | |
| MW581873.1 | | |
| MW581872.1 | | |
| MW581871.1 | | |
| MW581870.1 | | |
| MW581869.1 | | |
| MW581868.1 | | |
| MW581867.1 | | |
| MW581864.1 | | |
| MW581863.1 | | |
| MK370030.1 | | |
| MK286895.1 | | |
| MK086029.1 | | |
| MG280943.1 | | |
| MF580946.1 | | |
| KY575574.1 | | |
| KY575568.1 | | |
| KY038947.2 | | |
| KY038946.1 | | |
| KX262990.1 | | |

KU940225.1
KT324227.1
KP164569.1
KP164568.1
KP003809.1
KJ679577.1
JF274082.1
HQ456255.1
HQ456254.1
HQ456253.1
HQ456252.1
HQ456251.1
HM045823.1
HM045822.1
HM045821.1
HM045812.1
HM045811.1
HM045809.1
HM045805.1
HM045795.1
HM045794.1
HM045793.1
HM045792.1
HM045784.1
GU301781.1
GU301780.1
GU189061.1
GQ428211.1
FJ959103.1
FJ807899.1
FJ807898.1
FJ807896.1
FJ445510.2
FJ000069.1
FJ000068.1
FJ000067.1
FJ000066.1
FJ000065.1
FJ000064.1
FJ000063.1
FJ000062.1
EU564335.1
EU372006.1
EU244823.2
EU037962.1
EF210157.2
EF027139.1
EF027137.1

EF027136.1
EF027134.1
EF012359.1
DQ443544.2
AF490259.3
FN295487.2
LC259094.1
LC259093.1
AB455494.1
AB455493.1

Supplementary Table 2. CHIKV, ONNV, and MAYV primers and probes.

**CHIKV primers**

| | Forward Primer | | | | Reverse Primer | | | | |
|---|---|---|---|---|---|---|---|---|---|
| Target | sequence | size | GC% | binding site on NC_004162.2 | sequence | size | GC% | binding site on NC_004162.2 | Amplicon size |
| | CACCACCGACGTGATGAGACAGAGAGGTCTAG | 32-mer | 56% GC | Binds at: 2332 -> 2363 | CCAAGGCGATCAATGCAAGTAACGTTCCAG | 30-mer | 50% GC | Binds at: 2457 <- 2486 | 155 |
| | AATCACAACATCTGCACCCAAGTGTACCAC | 30-mer | 47% GC | Binds at: 2570 -> 2599 | CATCCTCTGAAGCACGTTAACACGAGATCTC | 31-mer | 48% GC | Binds at: 2739 <- 2769 | 200 |
| | GCCATTGTGTCATCGTTGCATTACGAAGGC | 30-mer | 50% GC | Binds at: 2636 -> 2665 | CCTCGTGTCCACGATAGTCAATTTGCAGTTG | 31-mer | 48% GC | Binds at: 2777 <- 2807 | 172 |
| | GAGTGGGAGGTGGAGCATGCATCAATAATGG | 31-mer | 52% GC | Binds at: 3026 -> 3056 | GGCTTGAATTATCTGGGACCACTGCCTGTCG | 31-mer | 55% GC | Binds at: 3166 <- 3196 | 171 |
| | ACGACAGGCAGTGGTCCCAGATAATTCAAGCC | 32-mer | 53% GC | Binds at: 3165 -> 3196 | CCGCGTAATACACAGACACCAACGGTTTAG | 30-mer | 50% GC | Binds at: 3288 <- 3317 | 153 |
| NSP2 | GAGCATATGGTTACGCAGATAGAACCAGTGAAC | 33-mer | 45% GC | Binds at: 3855 -> 3887 | GCTGCATTCAGTTGATTGTTCATGACATGAG | 31-mer | 42% GC | Binds at: 4014 <- 4044 | 190 |
| | CAACAGAAGTCAGTACGCCAGTCAGTGCTGC | 31-mer | 55% GC | Binds at: 5702 -> 5732 | TGCGCGACTGATACCTGCTTCTGTTGGCCATG | 32-mer | 56% GC | Binds at: 5836 <- 5867 | 166 |
| | CATTCCAGAACACACTACAGAATGTACTGG | 30-mer | 43% GC | Binds at: 6234 -> 6263 | AATTCTTCCCAGTATTCTTGGTTGCATGCG | 30-mer | 43% GC | Binds at: 6358 <- 6387 | 154 |
| | CTGCTGCCACTGCAGGAAGTGCCAATGGATAG | 32-mer | 56% GC | Binds at: 6488 -> 6519 | CCTAACCAGCTCTCTGTGAATCCCACATAGG | 31-mer | 52% GC | Binds at: 6631 <- 6661 | 174 |
| | GCTGTCATCTACCGACAGGTACGCGCTTCAAG | 32-mer | 56% GC | Binds at: 6903 -> 6934 | CCATGTATTATGTTGTCGTCGCCGATGAAG | 30-mer | 47% GC | Binds at: 7051 <- 7080 | 178 |
| NSP4 | TGCACGATATCGTGACAGGAACAGCTTGCAG | 31-mer | 52% GC | Binds at: 7197 -> 7227 | GTATACCGCTTTCTCCAACTCATCAATTAG | 30-mer | 40% GC | Binds at: 7349 <- 7378 | 182 |
| | CACAAGACCATACCTAGCTCACTGTCCCGAC | 31-mer | 55% GC | Binds at: 8574 -> 8604 | CGCAGCTTGGTCCAATCATGGCTATCATCC | 30-mer | 53% GC | Binds at: 8715 <- 8744 | 171 |
| | GCTACCTTGCAGCACGTACGTGCAGAGCAAC | 31-mer | 58% GC | Binds at: 8991 -> 9021 | GAGCCACCGCAATTACACTTATACCGCACC | 30-mer | 53% GC | Binds at: 9129 <- 9158 | 168 |
| E2 | CTACTATGACTGTAGTAGTTGTGTCAGTGG | 30-mer | 43% GC | Binds at: 9632 -> 9661 | GCTGTTCTGATGCAGCATATTAGGCTAAGCAG | 32-mer | 47% GC | Binds at: 9772 <- 9803 | 172 |
| | GAGTACCGTATAAGACTCTAGTCAACAGAC | 30-mer | 43% GC | Binds at: 10028 -> 1057 | GTATTCGCACGTGATGTAATCAAGCGATAG | 30-mer | 43% GC | Binds at: 10117 <- 10146 | 119 |
| | CGGCGACCATGCCGTCACAGTTAAGGACGC | 30-mer | 63% GC | Binds at: 10440 -> 10469 | GCTCTCAGGCGTGCGACTTTGGATATCGCC | 30-mer | 60% GC | Binds at: 10594 <- 10623 | 184 |
| E1 | ATCGACATACCGGATGCGGCCTTCACTAGG | 30-mer | 57% GC | Binds at: 10831 -> 10860 | CAGCTTCCCGGATAGTGACGGCGTTGGTCATC | 32-mer | 59% GC | Binds at: 10989 <- 11020 | 190 |

**CHIKV Probe**

| Target | sequence | Size | GC% | binding site on NC_004162.2 |
|---|---|---|---|---|
| E2 | /56-FAM/AGAAGGGCACTCGTGCCATAGT | 22-mer | 55% GC | Binds at: 8610->8631 |

**ONNV Primers**

| | Forward Primer | | | | Reverse Primer | | | | |
|---|---|---|---|---|---|---|---|---|---|
| Target | sequence | size | GC% | binding site on NC_075006.1 | sequence | size | GC% | binding site on NC_075006.1 | Amplicon size |
| | TGAAGAGTCATACGAACTGGTAAGGGTTGAG | 31-mer | 45% GC | Binds at: 2032 -> 2062 | ATATGGTGGGCTAGTTAAGTCGCCTACTAGC | 31-mer | 48% GC | Binds at: 2137 <- 2167 | 136 |
| | AACAGAACACGAGTACGTCTATGACGTCGATC | 32-mer | 47% GC | Binds at: 2065 -> 2096 | AGTCCTTCGTAGGCGAACTCATGATATGGTG | 31-mer | 48% GC | Binds at: 2160 <- 2190 | 126 |
| | GGTGTACACTGCCTGTAACAGCCATCGTGTC | 31-mer | 55% GC | Binds at: 2619 -> 2649 | CGGTAGTCTATTTGCAACTGCTTAACCCATC | 31-mer | 45% GC | Binds at: 2769 <- 2799 | 181 |
| | CGTTGCATTACGAGAGCAAGATGCGCACTAC | 31-mer | 52% GC | Binds at: 2652 -> 2682 | CTGTCATGACTTCGTTTCCACGGTAGTCTA | 30-mer | 47% GC | Binds at: 2790 <- 2819 | 168 |
| | ACTCCATTCCGCATACATCATTACCAGCAG | 30-mer | 47% GC | Binds at: 3746 -> 3775 | CTGGACGATCTGAACTTGCGTCCCAATACG | 30-mer | 53% GC | Binds at: 3901 <- 3930 | 185 |
| NSP2 | GTGTAGATCACGCAATGAAGCTCCAGATGC | 30-mer | 50% GC | Binds at: 3777 -> 3806 | CATTTCTGTATTGCTCGTGATGCACTGAGG | 30-mer | 47% GC | Binds at: 3941 <- 3970 | 194 |
| | CTCAGCTTCACCTGCCGCTGTGACACGATTG | 31-mer | 58% GC | Binds at: 887 -> 917 | TCACCGTCTACTGTATCTGTCGTCTTGCAC | 30-mer | 50% GC | Binds at: 1021 <- 1050 | 164 |
| | GTATCCTTCTCCGTGTGTACTTACGTACCAG | 31-mer | 48% GC | Binds at: 1055 -> 1085 | TTGCGTTGCGTCCTGCCATTGACCACTATC | 30-mer | 53% GC | Binds at: 1174 <- 1203 | 149 |
| | GTGTCCGAGAGAGGACCTTAACATGCTGTTG | 31-mer | 52% GC | Binds at: 1305 -> 1335 | ACCTGACGACCACAGGCTTGGTATCACGAAG | 31-mer | 55% GC | Binds at: 1426 <- 1456 | 152 |
| NSP1 | GGGCATTCAGAAAGCATAAGACGCATACGG | 30-mer | 50% GC | Binds at: 1341 -> 1370 | AGCCACTTGATTCTGGTTCTCAGCGGGATTG | 31-mer | 52% GC | Binds at: 1461 <- 1491 | 151 |
| | GCCATACACCTGTGGATGCAGACCGATCTGG | 31-mer | 58% GC | Binds at: 8854 -> 8884 | GCATGCATGTGTGACTGATCCTTCTACTGTCT | 32-mer | 47% GC | Binds at: 8987 <- 9018 | 165 |
| | GTTAAGATCACAGTTGACGGCCAGACAGTAC | 31-mer | 48% GC | Binds at: 9201 -> 9231 | GTCAACGGTGAGTTGTATTGCCATTTCTTGTG | 32-mer | 44% GC | Binds at: 9333 <- 9364 | 164 |
| E2 | GAGTCACTCTGCTGTTGTATCCAGACCATCC | 31-mer | 52% GC | Binds at: 9484 -> 9514 | CATGTGACCTCTAAGCCTTCTGCTGGTACT | 30-mer | 50% GC | Binds at: 9599 <- 9628 | 145 |
| | TACGAGCACGCAACAGTGATCCCGAACACG | 30-mer | 57% GC | Binds at: 10092 -> 10121 | CAATGTTGGTTCCAGAGTGACCGACTGTAG | 30-mer | 50% GC | Binds at: 10188 <- 10217 | 126 |
| | GGAGCAATACACGTGCCTTACTCCCAGGCAC | 31-mer | 58% GC | Binds at: 10770 -> 10800 | TGCATCTGGAATGTCAATGGAGACTGGTATG | 31-mer | 45% GC | Binds at: 10916 <- 10946 | 177 |
| E1 | CACTATCCGCGAACCTAACGTAGATGTCGAG | 31-mer | 52% GC | Binds at: 11102 -> 11132 | AGGAGGATGGCATGTCGCTGAGCAGTGTAC | 30-mer | 57% GC | Binds at: 11211 <- 11240 | 139 |

**ONNV Probe**

| Target | sequence | Size | GC% | binding site on NC_075006.1 |
|---|---|---|---|---|
| NSP2 | /56-FAM/GTTAAAGCCAGGAGGTTCACTTCTGATTAG/idSp/GCTTACGGGTACGCC/3SpC3/ | 30+15 | 51% GC | Binds at: 3829->3874 |

**MAYV primers**

| | Forward Primer | | | | Reverse Primer | | | | |
|---|---|---|---|---|---|---|---|---|---|
| Target | sequence | size | GC% | binding site on NC_003417.1 | sequence | size | GC% | binding site on NC_003417.1 | Amplicon size |
| | CCACCGAAAGACCACGTTGTTACATATCCAG | 31-mer | 48% GC | Binds at: 10972 -> 11002 | CACACAAGTAACTATTACCAGAATTAGCACGG | 32-mer | 41% GC | Binds at: 11099 <- 11130 | 159 |
| | GGCAGGTGCGACATCCATTCACATTCAAAC | 30-mer | 50% GC | Binds at: 10810 -> 10839 | CTGTGCACGTAGCACGCGAACTACAAACAG | 30-mer | 53% GC | Binds at: 10931 <- 10960 | 151 |
| | GGATGCATAATCCAGACAAATCCGGTCCGAG | 31-mer | 52% GC | Binds at: 10606 -> 10636 | GATGAGTGCGTGCATGTAGACACAGTGCAC | 30-mer | 53% GC | Binds at: 10734 <- 10763 | 158 |
| E1 | CACTCAACCTGGAGTATATCACTTGCGATTAC | 32-mer | 44% GC | Binds at: 9956 -> 9987 | GGCTTCGCTCATCTGTGTGTTCTCCGAATC | 30-mer | 53% GC | Binds at: 10123 <- 10152 | 197 |
| | GAGGCAGCATGACTTAGAGGTGCCGATTATG | 31-mer | 52% GC | Binds at: 5605 -> 5635 | GATTGGTATCTGCTCCGGTTGGCATCTGTTG | 31-mer | 52% GC | Binds at: 5721 <- 5751 | 147 |
| | CAGGGCAGCGCATATTACGTCTCTGCTGCG | 30-mer | 60% GC | Binds at: 5798 -> 5827 | CATCCGTAATCTGGTAGGACGCCACTGTAG | 30-mer | 53% GC | Binds at: 5955 <- 5984 | 187 |
| | GCCTCAATGCTGTGTTGCTTCCGAACATCC | 30-mer | 53% GC | Binds at: 6552 -> 6581 | GAAGCAATGTCCGTCTCCAGGACGTGGTCC | 30-mer | 60% GC | Binds at: 6646 <- 6675 | 124 |
| NSP4 | GTGGAGGTTTCCTGGTTTATGACCATGTTAC | 31-mer | 45% GC | Binds at: 7071 -> 7101 | GACCTCAATCTCAGAACCTAAGCCCGATCTAG | 32-mer | 50% GC | Binds at: 7227 <- 7258 | 188 |
| | CAGGCTTTGAGTGAGAGCGCAACGATGGTG | 30-mer | 57% GC | Binds at: 1931 -> 1960 | GTTTGCGGTCCACATCATAAACGTATTCGG | 30-mer | 47% GC | Binds at: 2073 <- 2102 | 172 |
| | CTGTCAGGAGATCATGCTTGATGTTAAGAG | 30-mer | 43% GC | Binds at: 2323 -> 2352 | GCATGGCATGCGAAAGCCTCATCGACGTAC | 30-mer | 57% GC | Binds at: 2431 <- 2460 | 138 |
| | CGTGGACACGAGGTCATGACAGCTGCTGCATC | 32-mer | 59% GC | Binds at: 2798 -> 2829 | CAGGGATCACCCGACAGCGTCTTCCACACAAG | 32-mer | 59% GC | Binds at: 2945 <- 2976 | 179 |
| | GCAACCTTGGAGGACTGGCAACGAGAGCATG | 31-mer | 58% GC | Binds at: 3017 -> 3047 | GGCCAAGATAATCGCACTCCAATCAGTGGC | 30-mer | 53% GC | Binds at: 3170 <- 3199 | 183 |
| | CCACGTGCTGTTGGTTAGTGAGTACAACCTC | 31-mer | 52% GC | Binds at: 3583 -> 3613 | CATTGTTGGTAGTGGTGTAGCCTATACGGTG | 31-mer | 48% GC | Binds at: 3744 <- 3774 | 192 |
| NSP2 | GAGCACCTACGCGTACGCTGACAGAACGAG | 30-mer | 60% GC | Binds at: 3853 -> 3882 | CAGGGTTACTGTCCTTCTACCATTGTCAAAG | 31-mer | 45% GC | Binds at: 3985 <- 4015 | 163 |
| | CACGCTCATCCTGGATATCGGCAGTGCACC | 30-mer | 60% GC | Binds at: 250 -> 279 | CTGTCAACGACTTCGCCTGATGCCTTAGCC | 30-mer | 57% GC | Binds at: 376 <- 405 | 156 |
| | GCGAAGATAGACGACCTGCAGTCAGTGATGG | 31-mer | 55% GC | Binds at: 416 -> 446 | GGTGCATGGACCGCATAGACGTCCTGATAG | 30-mer | 57% GC | Binds at: 523 <- 552 | 137 |
| NSP1 | GGTATGCGTGAACGTACCCTGACTTGCTGTTG | 32-mer | 53% GC | Binds at: 1301 -> 1332 | CGGTGTCTGATTCCTATCGACAACCCTGCG | 30-mer | 57% GC | Binds at: 1447 <- 1476 | 176 |

**MAYV Probe**

| Target | sequence | Size | GC% | binding site on NC_003417.1 |
|---|---|---|---|---|
| NSP2 | /56-FAM/CGGCGTTAAGCAGACTG | 17-mer | 58% GC | Binds at: 2404 -> 2420 |

Table 3. RT-RPA cross-reactivity testing.

| | RT-RPA tests | | |
|---|---|---|---|
| | CHIKV | ONNV | MAYV |
| CHIKV | + | - | - |
| ONNV | - | + | - |
| MAYV | - | - | + |
| Sindbis virus | - | - | - |
| Ross River virus | - | - | - |
| Dengue virus 1 | - | - | - |
| Dengue virus 2 | - | - | - |
| Dengue virus 3 | - | - | - |
| Dengue virus 4 | - | - | - |
| Zika virus | - | - | - |
| West Nile virus | - | - | - |
| Yellow fever virus | - | - | - |
| La Crosse virus | - | - | - |
| Snowshoe hare virus | - | - | - |
| SARS-CoV-2 | - | - | - |
